# IL-4 receptor alpha blockade in mice reduces skin inflammation, systemic response and the atopic march

**DOI:** 10.1101/2024.07.18.604194

**Authors:** Juan-Manuel Leyva-Castillo, Daniel Sen Hoi Wong, Raif S Geha

## Abstract

Atopic dermatitis (AD) commonly precedes food allergy and asthma in the atopic march. Epicutaneous (EC) sensitization in mice with ovalbumin (OVA) results in allergic skin inflammation that mimics AD and promotes food anaphylaxis or asthma following a gastric or intranasal challenge with OVA, respectively. Using our mouse model of EC sensitization with OVA we evaluate whether anti-IL-4Rα blocking antibody improve allergic skin inflammation and impact the atopic march. IL-4Rα blockade at the end of EC sensitization decreased allergic skin inflammation in OVA-sensitized mice, evidenced by significantly decreased eosinophil infiltration, decrease production of IL-4, IL-13 by OVA- restimulated splenocytes and reduced serum levels OVA-specific IgE. However, late IL- 4Rα blockade did not affect food anaphylaxis or airway inflammation in EC sensitized mice following an oral or intranasal challenge with OVA. IL-4Rα blockade at the beginning of EC sensitization not only impact allergic skin inflammation and systemic response to the antigen, but also decreased food anaphylaxis or airway inflammation following OVA challenge. These results suggest that the blockade of sensitization through the skin by IL-4Rα blockade could impact the atopic march.

## Introduction

Atopic dermatitis (AD) is characterized by a defective skin barrier function and a type 2-dominated local and systemic response to antigens encountered through the skin[1, 2]. AD commonly emerges in early life, followed in some individuals by the appearance of other allergic diseases such as food allergy, asthma, or rhinitis later in life, a progression named atopic march [3–6]. These results suggest that AD is the entry door for other allergic diseases.

We have previously shown that mice sensitized epicutaneously (EC), by the application of ovalbumin (OVA) on tape stripped skin, develop allergic skin inflammation with features of AD [7–10]. They also undergo IgE-mediated systemic anaphylaxis following a single oral antigen challenge or airway inflammation, and AHR with features of allergic asthma following intranasal challenge [8, 10–13]. This model mimics the atopic march observed in patients with AD, making this model an attractive tool to ascertain therapeutics to block the progression of the atopic march.

Genetic ablation experiments in mice have demonstrated the important role of IL- 4 and IL-13 in the development of allergic skin inflammation induced by EC sensitization [14–16]. In addition, local IL-4Rα blockade in EC sensitized mice with OVA decreased allergic skin inflammation and enhanced *S. aureus* clearance[17]. In patients with AD, systemic blockade of IL-4Rα, shared by the Th2 cytokines IL-4 and IL-13, improves the signs and symptoms of the disease, decreasing both the Th2 cells in circulation and total and allergen-specific IgE serum levels [18–22]. In addition, systemic blockade of IL-4Rα diminishes the appearance of new allergies [23]. These results suggest that IL- 4Rα blockade could be an interesting therapeutic option to control the atopic march. We made use of a monoclonal antibody against the murine IL-4Rα chain and our mouse model of atopic march induced by EC sensitization to investigate whether this therapeutic approach could control the progression of the atopic march.

## Materials and Methods

### Mice

BALB/c mice were purchased from Charles River Laboratory. All mice were kept in a pathogen-free environment and fed an OVA-free diet. All procedures were performed in accordance with the Animal Care and Use Committee of the Children’s Hospital Boston.

### Epicutaneous (EC) sensitization and antibody treatments

Female mice 6-8- weeks old were epicutaneously sensitized for 12 days. Briefly, Mice were anesthetized, and their back skin was shaved and tape-stripped with a film dressing (TegadermTM, 3M) followed by the application of 200 µg OVA (Sigma-Aldrich) or saline every other day. 100 μg of anti-IL-4Ra monoclonal antibody (clone mIL4R-M1, BD Biosciences), or IgG isotype control were intraperitoneal injected at different time points.

### Analysis of oral active anaphylaxis

Two weeks after the last sensitization, mice were challenged intragastrically with 150 mg of OVA in 350 μL of saline buffer. Temperature changes were measured every 5 minutes after OVA challenge by using the DAS-6001 Smart Probe and IPTT-300 transponders (Bio Medic Data Systems, Seaford, Del) injected subcutaneously. Sera were collected 60 minutes after challenge.

### Analysis of airway inflammation

Two weeks after the last sensitization, mice were challenged intranasally with OVA (50mg) daily for 3 days. Lung resistance was measured with invasive Buxco (Buxco Electronics, Wilmington, NC) in response to increasing doses of methacholine administered by means of nebulization to anesthetized mice 24 hours after the last intranasal treatment. Immediately after death, BALF and lung was collected for analysis.

### Determination of OVA-specific IgE in serum

Detection of OVA specific IgE was performed using a homemade sandwich ELISA as described previously [7, 17].

### Mouse serum mast cell protease 1 levels

Mouse mast cell protease 1 (mMCP- 1) concentrations were measured in sera collected 1 day before and 60 minutes after oral challenge by means of ELISA with a kit for mMCP-1 per the manufacturer’s instructions (eBioscience).

### Histology and measurement of epidermal thickness

Skin and lung specimens were fixed in 4% paraformaldehyde embedded in paraffin and H&E or PAS stained. ImageJ was used for the quantification of the epidermal thickness.

### Skin cell preparation

1cm^2^ skin pieces from EC sensitized mice were obtained. Skin pieces were finely chopped using scissors after fat removal and digested for 90 minutes in media containing Liberase (0.2mg/ml, Roche) and DNAse II (Sigma), with continuous shaking at 37° C. Digested skin homogenates were filtered, washed and resuspended in PBS and used for flow cytometry.

### Flow cytometry analysis

Single cell suspensions were preincubated with FcγR- specific blocking mAb (2.4G2) and washed before staining with the following monoclonal antibodies (mAbs): CD3 (17A2), CD45 (30F11), Gr1 (RB6-8C5) from eBioscience, CD11b (M1/70) and CD117 (2B8) from Biolegend and anti-Siglec-F (E50-2440) and anti-IgE (R35-72) from BD Biosciences. Cells were analyzed by flow cytometry using an LSR Fortessa machine (BD Biosciences). The data were analyzed with FlowJo software.

### RNA extraction and quantitative PCR analysis

RNA was extracted from whole skin or lungs with RNEasy mini kit (Qiagen). cDNA was prepared with iscript cDNA synthesis kit (Biorad). Quantitative real-time PCR was done using Taqman gene expression assays, universal PCR master mix and Quantstudio 5 Real Time PCR system (Applied Byosistems).

### Cytokines production by spleen cells

Spleen cell suspensions were cultured at 4×10^6^/ml in the presence of OVA (200 μg/ml) for 96 hours as described previously [7]. Cytokines in supernatants was measured by ELISA using Ready-Set-Go! ELISA Kits (eBioscience) following the manufacturer’s instructions.

### Statistical analysis

Statistical significance was determined by Student’s t test or one-way ANOVA analysis on Graph-pad prism. A p value <0.05 was considered statistically significant.

## Results

### Late IL-4Rα blockade reduces allergic skin inflammation and allergen sensitization induced by EC sensitization with OVA

The type 2 cytokines IL-4 and IL- 13 play an important role in allergic skin inflammation induced by EC sensitization with OVA [14–17]. We examined the effect of the systemic IL-4Rα blockade by monoclonal antibody in EC sensitized skin with OVA. Mice were EC sensitized with OVA for 10 days and intraperitoneally (*i.p.*) injected with anti-IL-4Rα blocking antibody or the IgG2a isotype control at day 7, 9 and 11, as illustrated in **Fig. 1A**. EC sensitization with OVA caused epidermal thickening, accumulation of T cells, mast cell and eosinophils, and local upregulation of *Il4*, *Il13*, and *Il17a* [7–10]. Late IL-4Rα blockade in EC sensitized skin with OVA did not affect epidermal hyperplasia or infiltration of CD45^+^ cells, however diminished eosinophil infiltration but not CD4^+^ T cells nor mast cells (**Fig. 1B-C**). In addition, IL-4Rα blockade did not affect the expression of cutaneous *Il4*, *Il13* or *Ifng* mRNA levels (**Fig. 1D**). However, EC sensitized skin *i.p.* injected with anti-IL-4Rα antibody exhibited an increased cutaneous *Il17a* mRNA level (**Fig. 1D**). Our results indicate that late IL-4Rα blockade decreases the infiltration of eosinophils and increase *Il17a* mRNA levels in EC sensitized skin with OVA.

**Figure 1.**
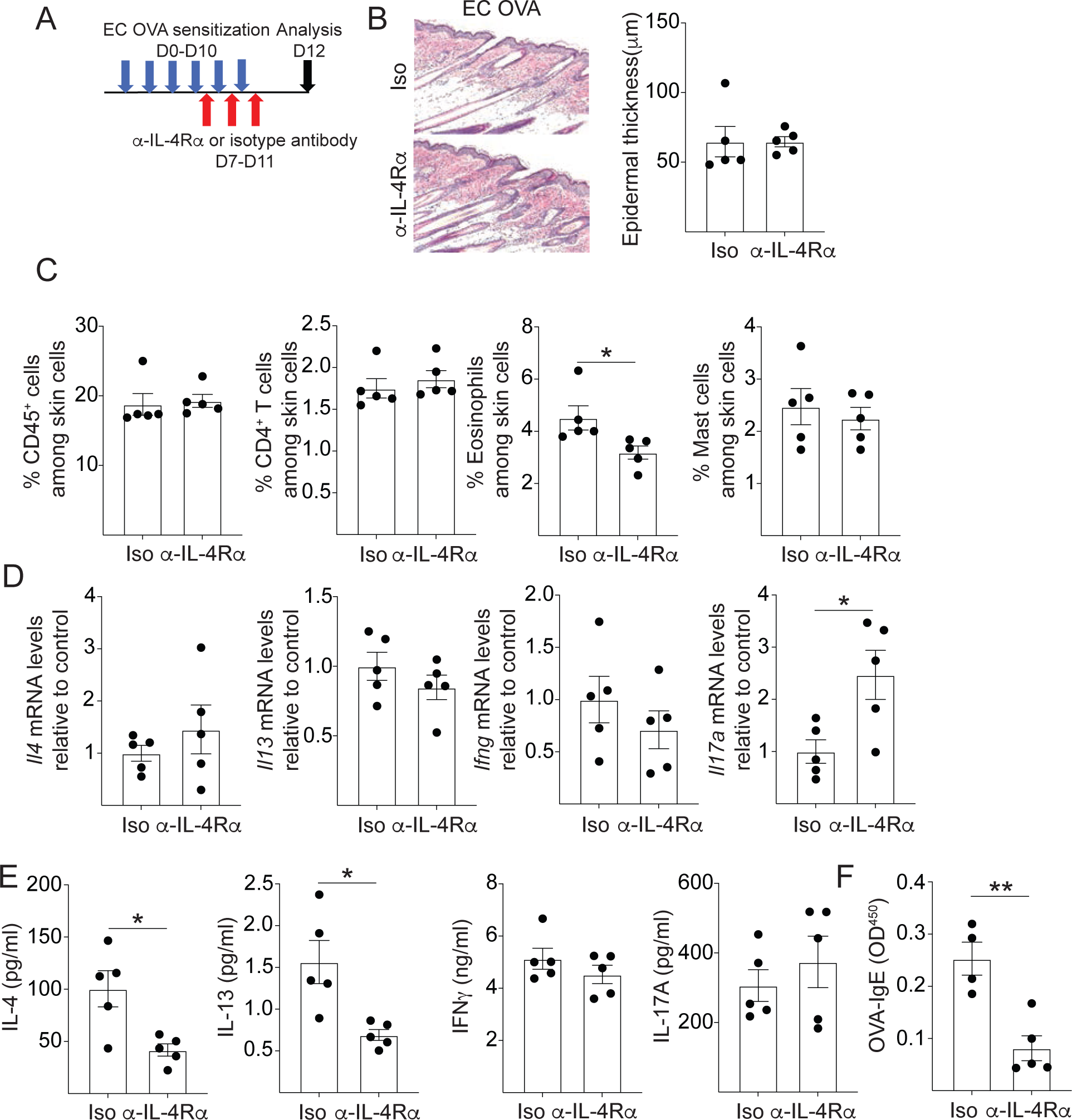
Late IL-4Rα blockade reduces eosinophil infiltration and systemic response induced by EC sensitization with OVA. A. Experimental protocol. **B-E.** Representative H&E staining and epidermal thickness (B), percentages of CD45^+^ cells, CD4^+^ T cells, eosinophils and mast cells (C) mRNA levels of cytokines expressed in the skin relative to the mean of isotype injected WT controls (D), IL-4, IL-13, IL-17A and IFNγ secretion by splenocytes (E) and serum OVA-specific IgE levels (F) in OVA-sensitized mice intraperitoneally injected with anti-IL-4Rα antibody or isotype control. Bars represent mean±SEM. * p<0.05, ** p<0.005.

EC sensitization with OVA promotes the secretion of splenocyte cytokines IL-4, IL- 13, IL-17A and IFNγ following OVA re-stimulation *in vitro* and the production of OVA-specific IgE antibodies [7–10]. Late IL-4Rα blockade in EC sensitized skin with OVA significantly reduced the secretion of IL-4, IL-13, but not IL-17A or IFNγ, by OVA restimulated splenocytes and serum OVA-specific IgE levels (**Fig 1E and F**). Our results indicate that late IL-4Rα blockade suppresses both cellular and humoral type 2 immune responses induced by EC sensitization with OVA.

### Late IL-4Rα blockade during allergic skin inflammation did not prevent food allergy or asthma in EC sensitized mice

We previously reported that oral or airway challenges using OVA in mice EC sensitized with OVA resulted in food anaphylaxis and asthma, respectively [8, 10–13]. To investigate whether the decrease in antigen specific immune response induced by systemic IL-4Rα blockade protects EC sensitized mice from food anaphylaxis or allergic asthma, we subjected EC OVA sensitized mice, *i.p.* injected with anti-IL-4Rα or isotype control, to oral or airway challenge with OVA (**Fig. 2A and 2D**). Following the oral challenge, EC OVA sensitized mice exhibit systemic anaphylaxis, characterized by a drop in body temperature and an increase in mMCPT-1 serum levels [11, 12]. Following the oral challenge, EC OVA sensitized mice injected with anti-IL-4Rα antibody *i.p.* exhibited comparable drop in body temperature and mMCPT-1 serum levels to EC OVA sensitized mice injected with isotype antibody (**Fig. 2B-C**).

**Figure 2.**
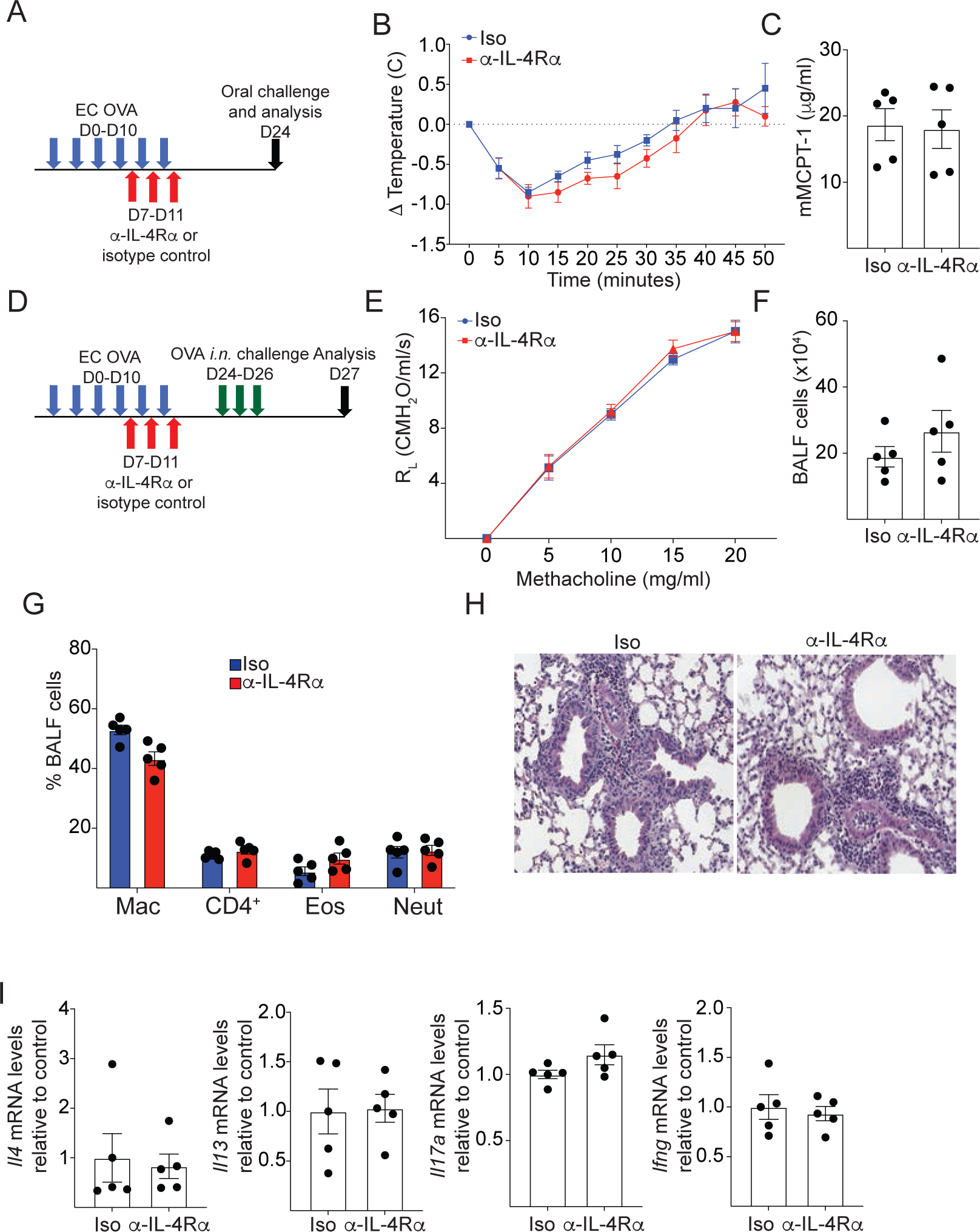
Late IL-4Rα blockade did not affect the progression of the atopic march in mice EC sensitized and challenged with OVA. A. Experimental protocol for food anaphylaxis. **B-C.** Change in body temperature (B) and serum mMCPT-1 levels (C) in OVA-sensitized mice intraperitoneally injected with anti-IL-4Rα antibody or isotype control. **D.** Experimental protocol for asthma. **E-I.** Lung resistance in response to increasing doses of methacholine (E) Total (F) and differential cell counts in BALF (G), H&E stained lung sections (H), and cytokine mRNA levels in lungs (I) of OVA- sensitized mice intraperitoneally injected with anti-IL-4Rα antibody or isotype control. Bars represent mean±SEM.

Intranasal OVA challenge in EC sensitized mice with OVA resulted in airway inflammation with features of allergic asthma and airway hyperresponsiveness (AHR) [8, 13, 16]. Following the intranasal challenge, anti-IL-4Rα antibody *i.p.* injected mice EC sensitized with OVA exhibited similar AHR, infiltration of total cell in bronchoalveolar lavage fluid (BALF) and comparable percentages CD4^+^ T cells, eosinophils, and neutrophils in to the BALF than isotype-treated controls (**Fig. 2E-G**). Lung histological analysis did not show any difference in the perivascular and peribronchial infiltration of mononuclear cells between the two groups (**Fig. 2H**). Moreover, *Il4*, *Il13*, *Il17a,* and *Ifng* mRNA levels in the lungs was comparable between the two groups (**Fig. 2I**). These results indicate that late IL-4Rα blockade during EC sensitization with OVA does not protect mice against experimental anaphylaxis or asthma following challenge with the same allergen.

### Early IL-4Rα blockade reduces allergic skin inflammation and allergen sensitization induced by EC sensitization with OVA

To investigate whether IL-4Rα blockade could be used as a preventive therapy for the development of allergic skin inflammation, mice were intraperitoneal injected with IL-4Rα blocking antibody or isotype control at the beginning of EC sensitization with OVA as displayed in figure **3A**. IL-4Rα blockade impairs the allergic skin inflammation induced by EC sensitization with OVA. Mice injected with IL-4Rα blocking antibody showed decreased epidermal thickness, reduced infiltration of immune cells (CD45^+^) in EC sensitized skin, including CD4^+^ T cells, eosinophils and mast cells compared with isotype injected controls (**Fig. 3B-C**). In addition, IL-4Rα blockade decreased *Il4* and *Il13* expression, but increased *Il17a* expression, without affecting *Ifng* mRNA levels in EC sensitized skin with OVA (**Fig. 3D**). These changes were accompanied by a decreased systemic allergen response, demonstrated by reduced secretion of IL-4 and IL-13, but not IL-17A or IFNγ, by splenocytes after *in vitro* OVA re-stimulation and decreased production of OVA-specific IgE (**Fig. 3E and F**). These results indicate that early blockade of IL-4Rα during EC sensitization in mice impact the development of allergic skin inflammation and antigen-specific systemic response.

**Figure 3.**
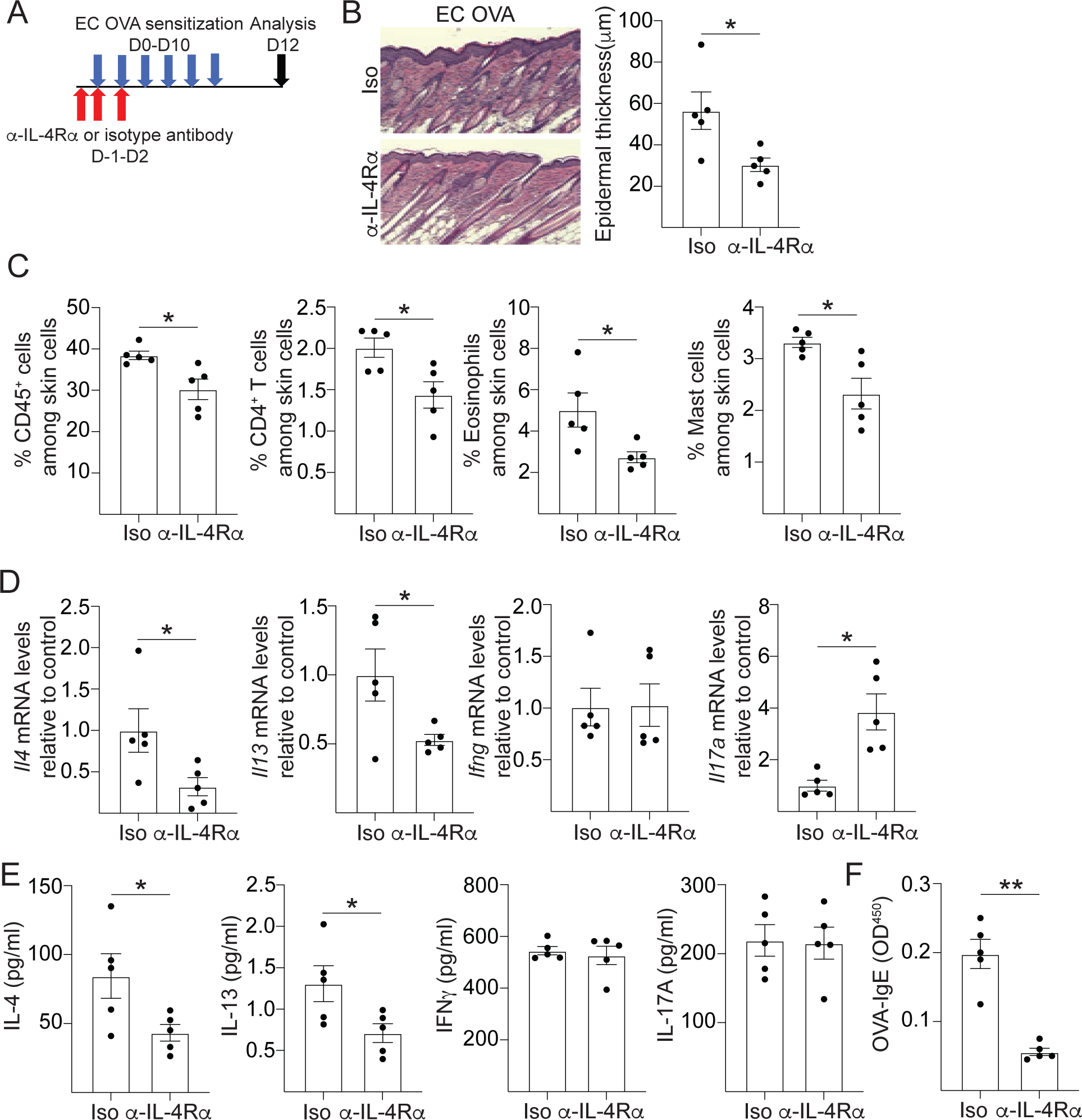
Early IL-4Rα blockade reduces allergic skin inflammation and systemic response induced by EC sensitization with OVA. A. Experimental protocol. **B-G.** Representative H&E staining and epidermal thickness (B), percentages of CD45^+^ cells, CD4^+^ T cells, eosinophils and mast cells (C) mRNA levels of cytokines expressed in the skin relative to the mean of isotype injected WT controls (D), IL-4, IL-13, IL-17A and IFNγ secretion by splenocytes (E) and serum OVA-specific IgE levels (F) in OVA- sensitized mice intraperitoneally injected with anti-IL-4Rα antibody or isotype control. Bars represent mean±SEM. * p<0.05, *** p<0.001.

### Early IL-4Rα blockade during allergic skin inflammation impact the progression of the atopic march

To investigate whether early IL-4Rα blockade prevents the progression to the atopic march, mice were intraperitoneal injected with IL-4Rα blocking antibody or isotype control at the beginning of EC sensitization with OVA, then these mice were orally or intranasally challenged with OVA as displayed in figure **4A** and **D**. Early IL-4Rα blocking antibody injection decreased the drop in core body temperature and serum mMCPT-1 levels in mice EC sensitized with OVA and orally challenged with OVA compared with those *i.p.* injected with isotype antibody (**Fig. 4B-C**). In addition, IL-4Rα blockade suppressed the AHR induced after intranasal challenge with OVA (**Fig. 4E**). This was accompanied by decreased infiltration of total cells in BALF and by decrease in the percentages of eosinophil infiltration in the BALF, decreased infiltration of mononuclear cells and mucus production in the lungs, and decreased expression of *Il4*, *Il13*, *Ccl24* and *Muc5a*, but not *Il17a* or *Ifng*, in the lungs (**Fig. 4F-I and data not shown**). Altogether these results indicate that early IL-4Rα blockade during the development of allergic sensitization protect mice against experimental anaphylaxis or asthma when the same allergen is re-encountered in another organ.

**Figure 4.**
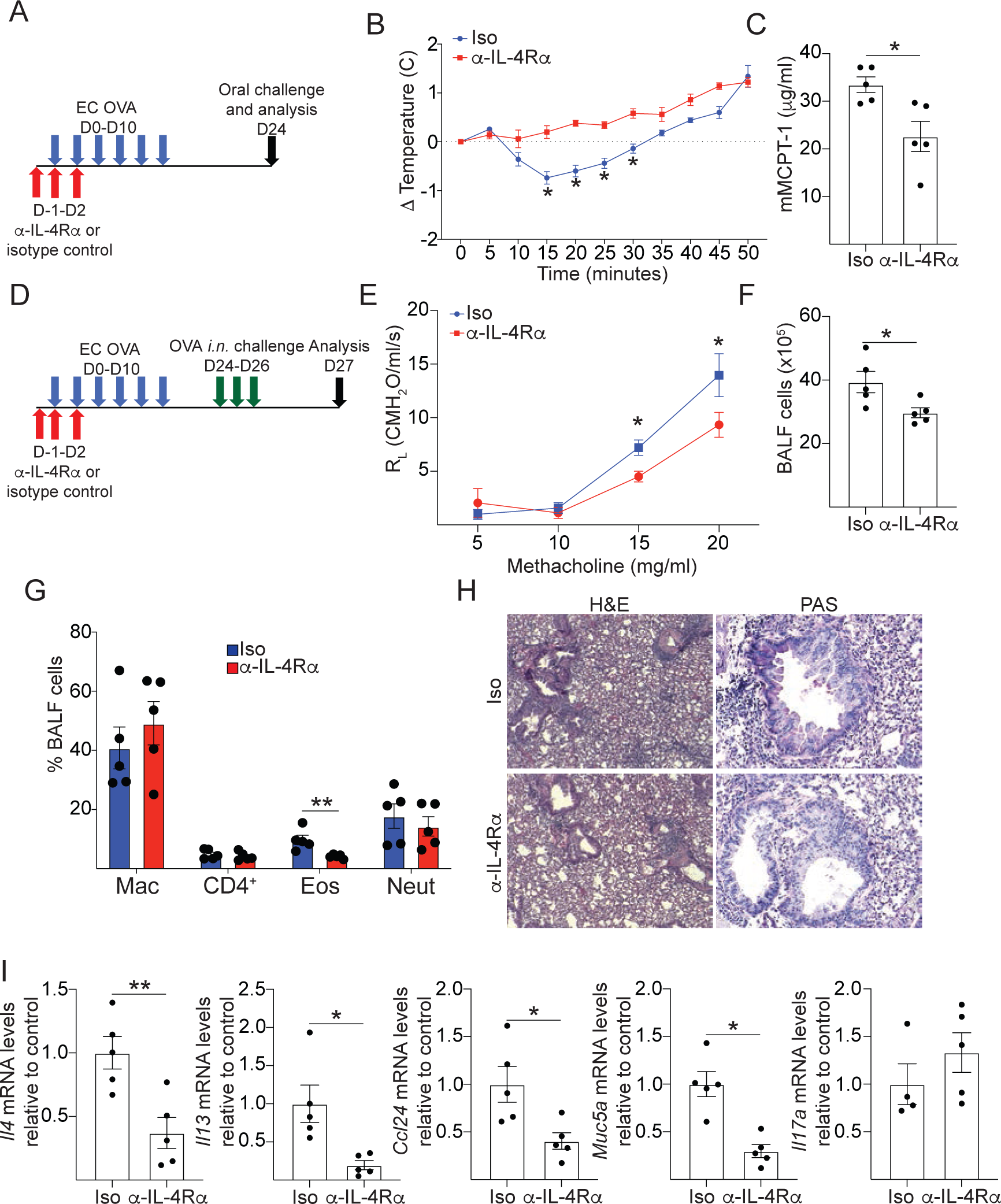
Early IL-4Rα blockade diminishes the atopic march in mice EC sensitized and challenged with OVA. A. Experimental protocol for food anaphylaxis. **B-C.** Change in body temperature (B) and serum mMCPT-1 levels (C) in OVA-sensitized mice intraperitoneally injected with anti-IL-4Rα antibody or isotype control. **D.** Experimental protocol for asthma. **E-I.** Lung resistance in response to increasing doses of methacholine (E) Total (F) and differential cell counts in BALF (G), H&E and PAS- stained lung sections (H), and mRNA levels in lungs (I) of OVA-sensitized mice intraperitoneally injected with anti-IL-4Rα antibody or isotype control. Bars represent mean±SEM. * p<0.05, ** p<0.005.

## DISCUSSION

Our results indicate that systemic IL-4Rα blockade at early or late time points of epicutaneous sensitization improves allergic skin inflammation, increases cutaneous Il17a mRNA levels and reduced allergen sensitization. However, only the early blockade of IL-4Rα improves food anaphylaxis and allergic airway inflammation following the reencounter of the allergen in the gastrointestinal or respiratory tract.

Late systemic IL-4Rα blockade decreased the accumulation of eosinophils in EC sensitized skin with OVA and decreased the systemic response against OVA. Previously we shown that a single intradermal injection with IL-4Rα blocking antibody in EC sensitized skin with OVA, in addition to reducing eosinophil, also decreased mast cells infiltration and epidermal thickness without affecting the systemic response to OVA. Altogether, these results suggest that IL-4Rα antibody concentration locally is important for its beneficial effect in the skin, while the concentration in circulation is important to decrease the systemic response against the allergen.

Our results showed that early and late IL-4Rα blockade increases cutaneous *Il17a* expression in EC sensitized skin. This is consistent with the role of IL-4 and IL-13 suppressing IL-17A expression in EC sensitized mouse skin and with recent reports indicating that dupilumab treatment in AD patients promote a psoriasis-like inflammation with increase in *IL17A* expression. [14, 17, 24–26].

Our results indicate that *i.p.* injection with IL-4Rα blocking antibody at early and later time points of EC sensitization with OVA impact the development of allergic skin inflammation and allergen sensitization. This is in line with the improvement of allergic skin inflammation in mice and lesional skin in AD patients, the decrease in allergen specific and total IgE and the decrease in Th2 cells evaluated by flow cytometry and single cell RNAseq observed in AD patients following IL-4Rα blockade [18-23, 27-30].

Recent reports indicating an important role of innate-derived IL-4 and IL-13 in the skewing of skin derived dendritic cells to a Th2 promoting phenotype during EC sensitization [31, 32]. In line with this, our results indicate that IL-4Rα blockade at the initiation, but not at the end, of EC sensitization with OVA, diminished the experimental allergic asthma and food anaphylaxis following intranasal and oral challenges with OVA, respectively. Our results are consistent with recent report indicating that treatment with Dupilumab in highly atopic patients with AD decreases sensitization to new allergens and prevents the appearance of new allergic conditions[23]. Altogether these results strongly suggest that the potential preventive role of IL-4Rα blockade in the progression of the atopic march is due to the inhibition of allergen sensitization encountered through a defective skin barrier.

## Abbreviations

AD: Atopic dermatitis
AHR: airway hyperresponsiveness
BALF: bronchoalveolar lavage fluid
EC: Epicutaneous
*i.p.*: intraperitoneally
mMCP-1: Mouse mast cell protease 1
OVA: Ovalbumin

